# Enterotoxigenic *Escherichia coli* display a distinct growth phase before entry into stationary phase with shifts in tryptophan- fucose- and putrescine metabolism and degradation of neurotransmitter precursors

**DOI:** 10.1101/2021.08.24.457600

**Authors:** Enrique Joffré, Xue Xiao, Mário S. P. Correia, Intawat Nookaew, Samantha Sasse, Daniel Globisch, Baoli Zhu, Åsa Sjöling

## Abstract

Enterotoxigenic *Escherichia coli* (ETEC) is a major cause of diarrhea in children and adults in endemic areas. Gene regulation of ETEC during growth *in vitro* and *in vivo* needs to be further evaluated, and here we describe the full transcriptome and metabolome of ETEC during growth from mid-logarithmic growth to stationary phase in rich medium (LB medium). We identified specific genes and pathways subjected to rapid transient alterations in gene expression and metabolite production during the transition between logarithmic to stationary growth. The transient phase during late exponential growth is different from the subsequent induction of stationary phase-induced genes, including stress and survival responses as described earlier. The transient phase was characterized by the repression of genes and metabolites involved in organic substance transport. Genes involved in fucose and putrescine metabolism were upregulated, and genes involved in iron transport were repressed. Expression of toxins and colonization factors were not changed, suggesting retained virulence. Metabolomic analyses showed that the transient phase was characterized by a drop of intracellular amino acids, e.g., L-tyrosine, L-tryptophan, L-phenylalanine, L-leucine, and L-glutamic acid, followed by increased levels at induction of stationary phase. A pathway enrichment analysis of the entire transcriptome and metabolome showed activation of pathways involved in the degradation of neurotransmitters aminobutyrate (GABA) and precursors of 5-hydroxytryptamine (serotonin). This work provides a comprehensive framework for further studies on transcriptional and metabolic regulation in pathogenic *E. coli.*

**Importance:** We show that *E. coli*, exemplified by the pathogenic subspecies enterotoxigenic *E. coli* (ETEC), undergoes a stepwise transcriptional and metabolic transition into the stationary phase. At a specific entry point, *E. coli* induces activation and repression of specific pathways. This leads to a rapid decrease of intracellular levels of L-tyrosine, L-tryptophan, L-phenylalanine, L-leucine, and L-glutamic acid due to metabolism into secondary compounds. The resulting metabolic activity leads to an intense but short peak of indole production, suggesting that this is the previously described “indole peak,” rapid decrease of intermediate molecules of bacterial neurotransmitters, increased putrescine and fucose uptake, increased glutathione levels, and decreased iron uptake. This specific transient shift in gene expression and metabolomics is short-lived and disappears when bacteria enter the stationary phase. We suggest it mainly prepares bacteria for ceased growth, but the pathways involved suggest that this transient phase substantially influences survival and virulence.

## Background

*Escherichia coli* is a facultative anaerobic gram-negative bacterium that normally inhabits the intestines of mammals and reptiles as a commensal bacterium. Pathogenic *E. coli* have acquired extrachromosomal genetic properties that enable them to colonize and adhere to the epithelium, thereby delivering toxins or virulence factors that harm the host [1]. The virulence factors can either be located on plasmids or inserted in the chromosomes as pathogenicity islands, and pathogenic *E. coli* can be found in most *E. coli* phylogroups [2–4]. Enterotoxigenic *Escherichia coli* (ETEC) is characterized by the production of the heat-labile toxin (LT) and/or the heat-stable toxin (ST). In most cases, each bacterium expresses one to three colonization factors that mediate adhesion to the epithelium in the small intestine [5, 6]. The toxins and colonization factors of ETEC are mainly encoded on extrachromosomal plasmids that have been acquired by horizontal transfer to ancestral commensal *E. coli*. Successful combinations of host and plasmid may lead to global transmission of virulent clones with optimal colonization and survival abilities [2, 4].

The molecular events that govern the virulence of ETEC are less well characterized than several other intestinal pathogenic *E. coli*. Studies on ETEC have so far focused on a few global regulators such as Crp, H-NS, and members of the AraC family such as Rns and CsvR that were shown to regulate the toxins and/or CFs [7–11]. However, virulence factors are not only the expression of toxins or colonization factors but can also be expanded to involve the ability to persist host-induced stress or to facilitate spreading or colonization ability. Studies using real-time PCR, microarrays, and RNA-Seq have recently begun to elucidate the global and/or specific transcriptional regulation in ETEC in response to host environmental factors such as bile and glucose and adhesion to epithelial cells [8, 11–15].

In this study, we explored the transcriptional and metabolomic profile of two ETEC clinical isolates belonging to a globally spread linage during growth from logarithmic cell division to early stationary phase to elucidate the effect of growth on transcriptomic profile and virulence gene regulation. We used a multi-omics approach including RNA-seq and mass spectrometric global metabolomics techniques to analyze the global regulation and provide a framework of *E. coli* genes, transcription factors, and analysis of metabolites involved in different growth phases in Luria Bertani medium.

## Results

### Expression profiling of two ETEC revealed characteristic transcriptional patterns during transition from log phase to early stationary phase

To profile the ETEC transcriptome during bacterial growth transition from mid-exponential to early stationary phase, we performed RNA-seq analysis on RNA isolated from two clinical isolates of ETEC (E1777 and E2265) grown in LB media. We sequenced the transcripts expressed after 3, 4 and 5 hours of growth corresponding to mid-log phase (3h, OD_600_ = 1.1-1.3), late log phase (4h, OD_600_ = 2.8-3.1) and early stationary phase (5h, OD_600_ = 4.6-4.8) (Figure 1).

**Figure 1.**
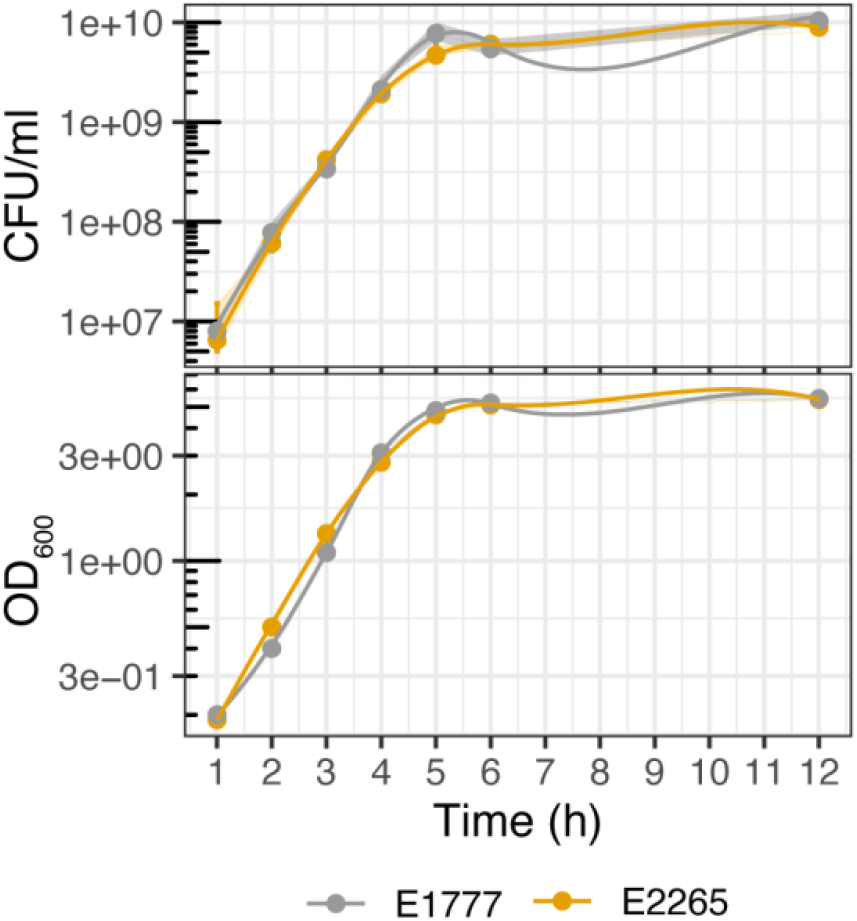
Growth curve of ETEC strains. Total bacterial counts in colony-forming units (CFU/ml) and optic density (OD_600_) of the ETEC strains E2265 and E1777 in LB media. Samples were measured every hour for 12 hours.

The two strains express LT STh, CS5+CS6, and belong to the globally distributed L5 ETEC lineage [4]. The time points were chosen to reflect the transition from active growth to carbon and nutrient starvation, an environment that enteropathogens face in the human gut and growth in LB [16]. First, Illumina sequence reads were assembled and assigned as unigene sequences, translated, and compared to protein databases for annotation. As a result, a total of 4166 unigenes with expression in at least one sample were detected. Next, in order to identify the differentially expressed genes shared by both isolates E2265 and E1777 and display the dynamic of the transcriptome during the transition from the log phase (3h) to the early stationary phase (5h), we performed a differential gene expression analysis by DESeq with a fold-change cutoff of 4 (Log2>2) and p-value < 0.001. Overall, 617 genes (S Table 1, S Table 2) showed a significant change (up/down) in at least one-time point, and these results were displayed and clustered in a heatmap in Figure 2. The following comparisons, 3 h *vs*. 4 h and 3 h *vs*. 5 h resulted in a number of 486 (257 upregulated and 229 downregulated) and 392 (209 upregulated and 183 downregulated) differently expressed genes (DEG), respectively (S Table 2). Thus, a total of 495 filtrated differentially expressed genes (DEG) in at least one condition in both bacterial transcriptomes were included to perform the *K*-means clustering analysis of the DEG heatmap (Figure 2 a; S Table 3) resulting in four clusters of specific gene expression patterns (Cluster I-IV). In parallel, the Short Time-series Expression Miner (STEM) clustering method [17] was performed using the same dataset to identify differential gene expression patterns shared among genes with similar dynamics in transcriptional changes over time (S Figure 1).

**Figure 2.**
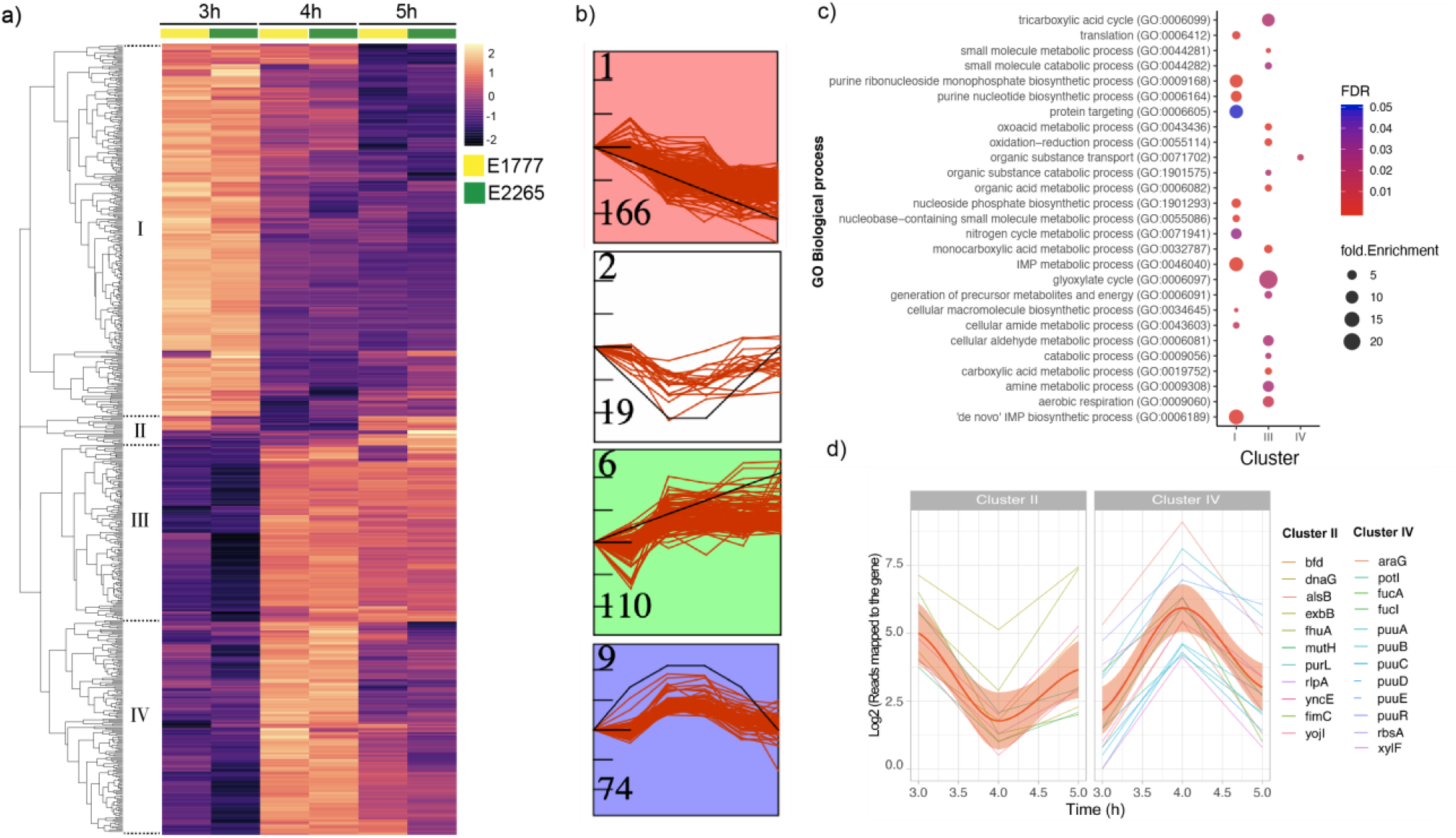
Transcriptomic response of ETEC during bacterial growth transition from mid-exponential to early stationary phase. A) Heatmap of the differential expressions of the two strains E1777 and E2265 after 3, 4, and 5 hours of growth in LB medium. K-*mers* analysis indicated 4 main clusters (I-IV). B) STEM analysis identifying the most common patterns of gene expression. C) GO Biological enrichment analysis of significant genes per cluster and fold enrichment. D) Set of genes with transient up or down-regulation.

Thus, STEM (Figure 2 b and S Figure 1) identified eight temporal expression profiles, of which four enclosed the majority of the genes, of which three (labeled in red, green, and blue) showed a statistically significant (*p* < 0.05) higher number of genes assigned using a permutation test. Both methods resulted in four distinctive temporal dynamics of the transcriptome in response to the transition from log to early stationary phase (S Table 4). In summary, clusters 1 and II were the largest, enclosing 166 and 110 genes, respectively. Cluster I showed a decreasing expression towards the entry to the stationary phase (3 h – 4 h – 5 h), while cluster III showed the opposite trend. Interestingly, clusters II and IV included 19 and 74 genes, respectively, with a significant transient down or up-regulation at 4 hours compared to 3 and 5 hours. Thus, the data indicate that a specific transient phase in gene regulation occurs after 4 hours and OD_600_ around 3 when ETEC enters the late log phase and starts transit into the stationary phase.

In order to gain biological insights from the temporal transcriptomic dynamics, we performed gene ontology (GO) enrichment analysis for biological processes and metabolic pathways of the significantly expressed genes from each cluster (Figure 2 c; S Table 5). Significant gene enrichment (FDR < 0.05) of cluster I indicated a notorious downregulation of genes involved in the *de novo* purine biosynthesis pathways such as ‘de novo’ IMP biosynthetic processes as well as nitrogen metabolic pathways, and protein targeting (intracellular protein transport) when growth slowed down upon entry to late log phase and early stationary phase. In contrast, cluster III, which represents a progressive transcriptional gene activation towards the stationary phase, was significantly enriched in glyoxylate metabolism, TCA cycle, carboxylic acid metabolic processes, aerobic respiration, and small molecule metabolism. Organic substance transport, which involves the movement of organic substances that contains carbon in, out, or within a cell, was solely identified as the most enriched biological process among genes of cluster IV. No significant enrichment was identified for Cluster II. Even though Clusters II and IV represent a minor proportion of the DEG dataset, they exhibited an interesting transient transcriptional response prior to switching into the stationary phase.

### Transient transcriptomic activation of putrescine and fucose utilization and reduction of iron transport prior to entry into stationary phase

The next step is to increase our understanding of the biological role of the transiently altered genes (4 hours of growth at the beginning of the stationary phase) identified in clusters II and IV. Reads mapped to the gene of each temporal gene expression were plotted in Figure 2d. The trendline confirmed the down (cluster II) and up (cluster IV) regulation of these genes during the transient shift to stationary phase. Among the activated genes at 4 h, we identified an overrepresentation of genes from the fuc operon, *i.e*., *fucI* (L-fucose isomerase) and *fucA* (L-fucose 1-phosphate aldolase) and *fucU* (L-fucus mutarotase) with an approximately 6-fold increase in expression. Another set of genes involved in the exploitation of alternative nutrient sources was the putrescine pathway. Like the fucose operon, the Puu-operon (putrescine utilization pathway) genes that degrade putrescine to GABA via γ-glutamylated intermediates were highly activated at 4 h. Genes involved in the putrescine uptake system, *i.e*., *potI* and *ydcU* showed the same pattern. Another gene included in this cluster was *tnaA*, which encodes tryptophanase, responsible for indole production from L-tryptophan. Its expression was activated 20-fold at 4 h compared to the mid-log growth phase at 3 h and reduced 6-fold at 5h (S Table 3). In addition, increased expression of the fad-operon (*fadH, FadM*) and the *dpp*-operon in charge of dipeptide transport were evident.

Downregulation of genes in cluster II included *exbB* and *fhuA* involved in siderophore-mediated iron transport, *yncE*, a DNA binding protein involved in iron metabolism, and *bfd* bacterioferritin-associated ferredoxin; hence cluster II suggests that a rapid downregulation of genes involved in iron metabolism occurs transiently before entry into early stationary phase.

### Expression of ETEC virulence factors is slightly higher during exponential growth

Both strains analyzed in this study expressed genes encoding the enterotoxins: *eltAB* (LT) and *estA* (STh), and colonization factor operons: *csfABCDEF* (CS5) and *cssABCD* (CS6) [18]. The transcriptome data showed that *eltAB* expression was higher than *estA* at all three-time points. In comparison with *estA* expression, *eltAB* was 3-4-fold higher at 3 hours and 2-4-fold and 2-fold at 4 and 5 hours, respectively. Expression of *eltAB* and the CS5 encoding *cfs*-operon, *estA*, and the CS6 had a trend of gradual downregulation over time (S Table 4). The expression patterns were similar between the strains. Although the expression levels were not significantly changed in this study, the findings are confirmed by our previous work [8]. Genome analysis showed that both strains also express additional virulence genes: *cexE, clyA, eatA, ecpA*, and *fimH* [18]. These genes were not significantly changed.

### Global view of the intracellular and secreted metabolome of ETEC growth phases

Since our transcriptomic data provided a framework of how metabolic pathways were altered during the transition from freely available nutrients to a more restricted environment, we wanted to characterize the impact of nutrient depletion on the bacterial metabolome. We collected pellets (intracellular metabolites) and supernatants (secreted metabolites) of E2265 and E1777 bacteria at 3, 4, and 5 hours of growth in LB broth medium and performed a nontargeted GC-MS-based metabolomics approach. Approximately 2000 putative intracellular and secreted metabolites were detected in both isolates, of which 288 metabolites were successfully identified (S Table 6). Principal component analysis (PCA) (Figure 3a) of all samples revealed a profound clustering of samples from the intracellular and secreted metabolomes. The analysis also indicated that the intracellular metabolomes of both E2265 and E1777 are more similar than their respective secreted metabolome. The PCA did not show any variation between metabolomes of the strains per time point (Figure 3a); however, the hierarchical clustering heatmap showed changes in the abundance of several metabolites along with the time points (S Figure 2).

**Figure 3.**
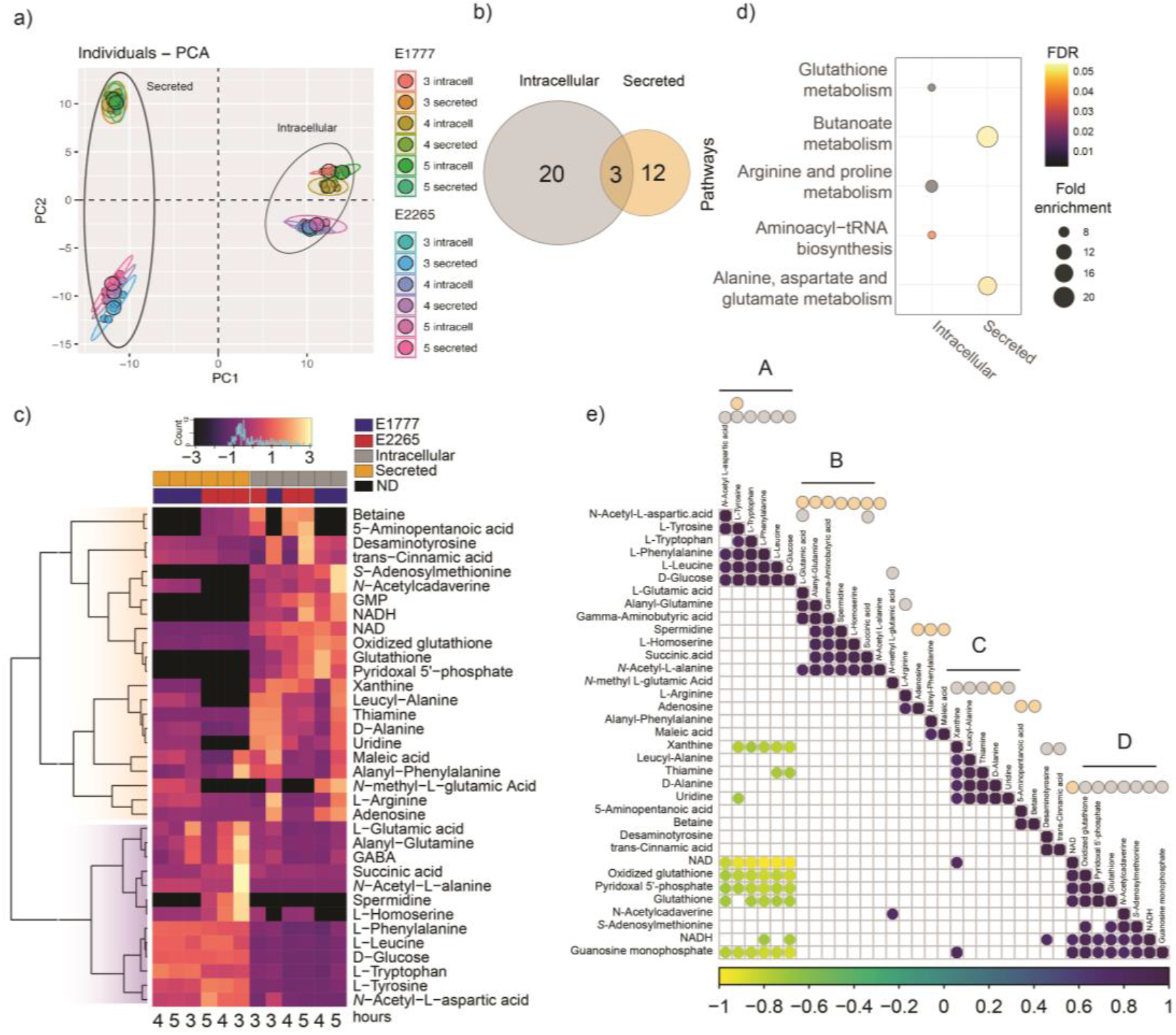
Intracellular and Secreted profile of the metabolic response to ETEC during bacterial growth transition from mid-exponential to early stationary phase. A) PCA plot generated from all metabolites of different samples. B) Venn diagrams of number of significant (p < 0.01 −2 > Log2Foldchange <2) found in the bacteria (intracellular) or medium (secreted). C) Heatmap representation of the 35 differentially changed metabolites at any time point. D) Pathway enrichment analysis of the intracellular and secreted significant metabolites. Non-significant pathways were colored gray. E) Metabolite-metabolite correlation analysis shows positive correlations in dark purple and negative correlations in yellow. Secreted and intracellular metabolites were marked with orange and gray dots, respectively.

#### Intracellular and secreted metabolome show unique metabolomic shifts

Since we are interested in studying the differences in the metabolome during bacterial growth transition mid-exponential to early stationary phase, we used multiple t-tests to compare the metabolite abundance between two time points, *i.e*., 3 h versus 4 h or 3 h versus 5 h (S Table 7, S Table 8). The threshold for significant changes was |log_2_ fold change > log_2_ 2; Padj < 0.05). As shown in Table 1, Figure 3 b, and S Figure 3, 35 metabolites significantly changed in their intracellular and/or extracellular concentration at any time point. Specifically, 20 intracellular and 12 secreted metabolites were significantly altered.

**Table 1.**
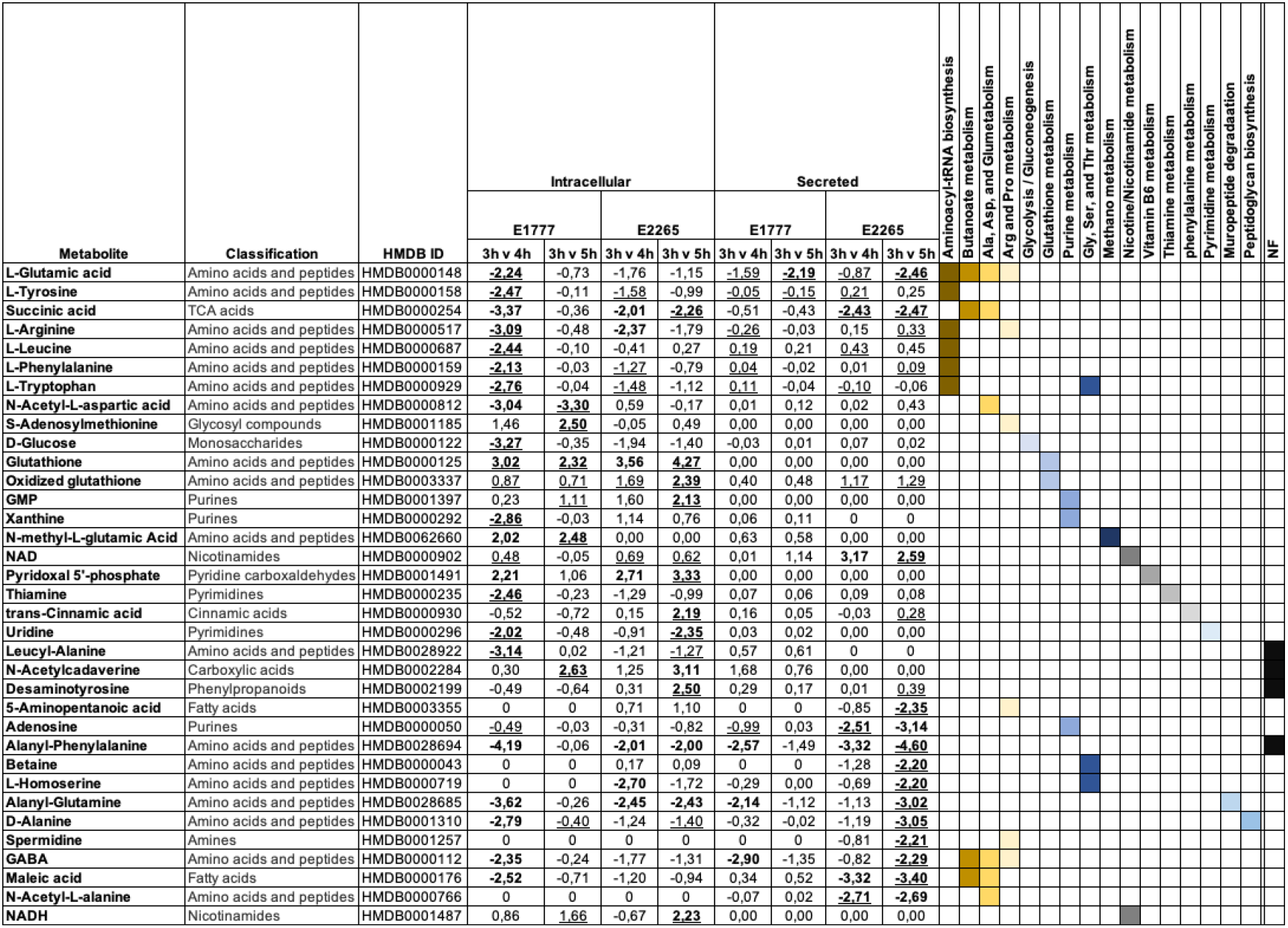
Summary of significantly altered intracellular and secreted metabolites of ETEC. Metabolites were classified based on the biochemical structure and in which metabolic pathways play a role. Metabolite abundance is presented in fold change against 3 h. Values in bold represent −2 > Log2Foldchange < 2 and underlined were statistically significant Padj < 0.05.

To characterize the chemical diversity of the intracellular and secreted ETEC metabolomes and further explore the metabolic changes between the intracellular and secreted ETEC metabolomes across the time points, the significant metabolites were classified according to chemical classes (Human Metabolome DB; www.hmdb.ca) and a hierarchical clustering heatmap based on the metabolite relative abundances was generated. Ten and five metabolite classes were included in the intracellular and secreted metabolome, respectively. The most common class of metabolites were amino acids and peptides, as well as downstream catabolism metabolites. The metabolic profile changes illustrated in the heatmap of Figure 3c indicated remarkable differences in the metabolite abundance between ETEC metabolomes and some differences in the metabolic profile between ETEC strains. For instance, betaine and the fatty acid 5-aminopectanoic acid were only detected in E2265, and the amino acid *N*-methyl-L-glutamic acid only in E1777 (Figure 3c).

Two main clusters of metabolite abundance patterns were identified: the first cluster included a diverse set of metabolites with higher intracellular concentrations or absence of secretion. For instance, *S-*adenosylmethionine, *N-*acetylcadaverine, GMP, NADH, glutathione, and pyridoxal 5′-phosphate (PLP, known as the catalytically active form of vitamin B_6_) were not secreted at any time point by the two strains.

In the second cluster with higher levels of secreted metabolites than intracellular levels, we found most of the amino acids identified in this dataset and glucose and succinic acid. The amine spermidine was only detected in the E2265 supernatant, whereas the amino acid L-homoserine was secreted at higher levels by E2265 than E1777, where the metabolite was absent intracellularly (Figure 3c). Again, these data indicated that although both ETEC strains are genetically very closely related, their physiology could vary. The clustering patterns of samples per time point also confirmed significant differences in metabolites abundance over time.

A pathway enrichment analysis of significantly altered metabolites discovered general metabolic differences between both fractions. As is shown in Figure 3d, three KEGG pathways were differentially enriched (FDR < 0.05), and several other pathways were associated with the significant metabolites (Table 1). The intracellular metabolites with significant differences over the timepoints were enriched in the aminoacyl-tRNA biosynthesis pathway, including L-tyrosine, L-tryptophan, L-phenylalanine, L-leucine, and L-glutamic acid. These metabolites showed significantly decreased levels at 4 hours which were restored in E1777 at 5 hours but remained low in E2265. The intracellular glutathione levels increased 10-fold at 4 h compared to 3 h and remained high at 5 h. On the other hand, the pathways responsible for butanoate metabolism, alanine, aspartate, and glutamate metabolism were significantly enriched among secreted metabolites (Figure 4 c and S Table 9).

**Figure 4.**
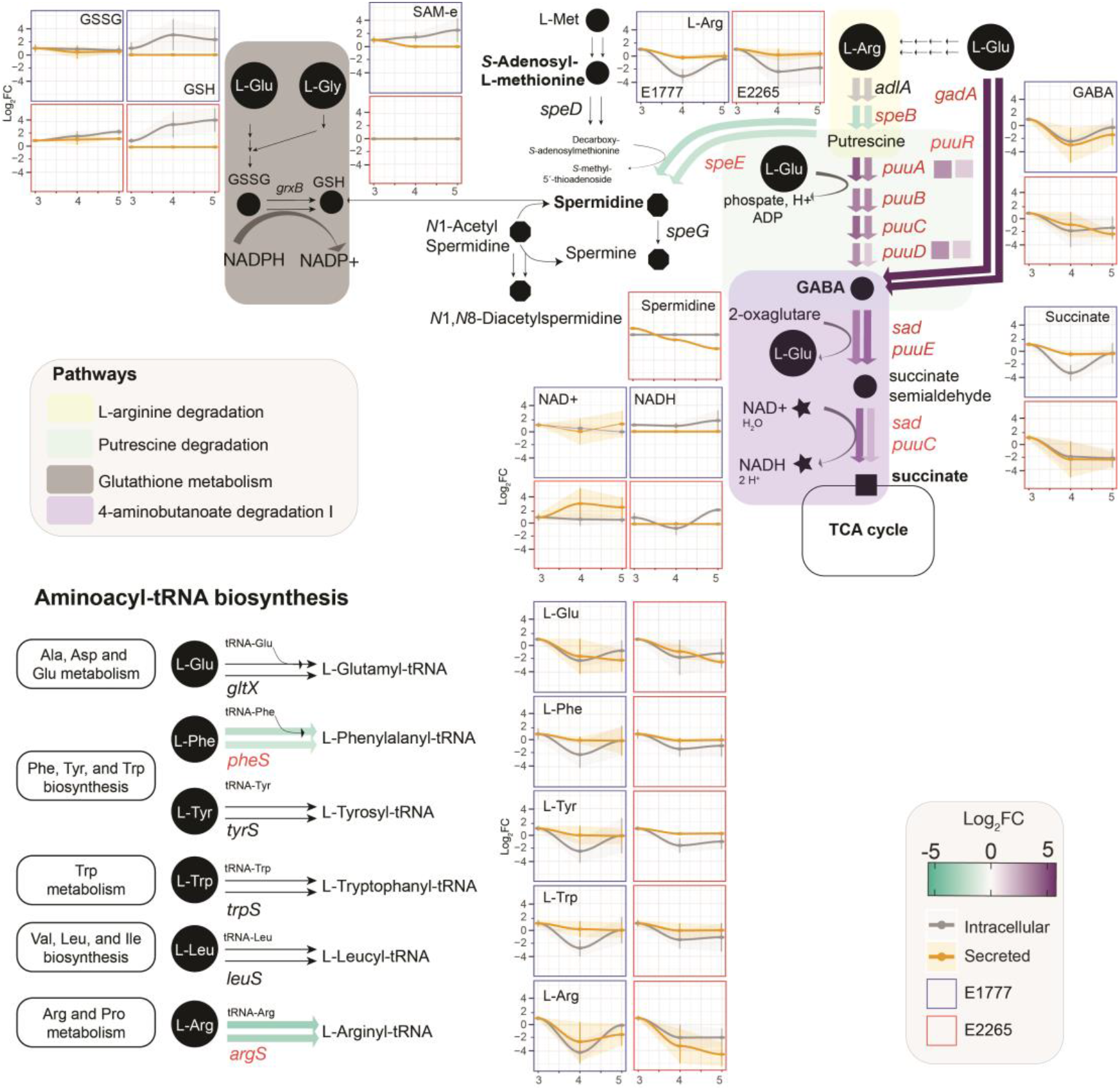
Transcriptomic and metabolic modulation of 4-butanoate degradation, related pathways, and Aminoacyl-tRNA biosynthesis of ETEC during growth. Circles are metabolites, and arrows are reactions. Verified metabolites are labeled in bold (S Table 12). The color of the width arrows indicates the levels of gene expression represented in the fold-change. The first and second arrows represent the DEG between 3 vs. 4 h and 3 vs. 5h. Data of the concentration of significant metabolites are represented in the serial charts. The metabolite concentration is adjusted to 3 h, and the data is presented in fold-change±SD.

The butanoate metabolism pathway included the bacterial neurotransmitter GABA, L-glutamate, succinic acid, and maleic acid, and they were mainly secreted at 3 hours. These metabolites were found in lower concentrations in comparison with metabolites of the aminoacyl-tRNA biosynthesis pathway.

In order to investigate whether there were metabolite-metabolite correlations across the metabolomes, we calculated the Pearson correlation coefficients. We wanted to identify the strongest correlations, and a total of 110 pairwise differentially significant correlations (P < 0.001) were found, of which 74 were positive and 36 negative metabolite-metabolite correlations. Most of these correlations were consistent with the heatmap clustering; however, 4 sets of tightly correlated metabolite-metabolite interaction were identified. As seen in Figure 3 e (S Table 10), all 4 grouped pairwise metabolites positively correlated (r < 0.8). Most of the amino acid metabolites were clustered in groups A and B, while the C and D groups included more diverse classes of metabolites. The positive correlations emphasized their close biochemical relatedness or overlapped roles among catabolic pathways. In contrast, metabolite-metabolite correlations from A and D as well as xanthine, thiamine, and uridine from the nucleotide metabolisms and A correlated negatively (r < −0.6).

Overall, the metabolome analysis of ETEC revealed distinctive differences in the metabolic composition of the intracellular and external environment of the bacteria, mainly characterized by the secretion or availability of large amounts of essential amino acids, intermediates, and derived molecules such as GABA. In contrast, the intracellular environment was characterized by a transient drop of amino acids at 4 hours of growth, and glutathione redox-regulation was implicated in the adaptation to stationary phase as reduced L-glutathione was increased 10-fold intracellularly.

### Transcriptome and metabolome integration

Next, we combined the significant data from the transcriptomic analysis and metabolomics and mapped it into super pathways, including L-arginine, 4-aminobutanoate, and putrescine degradation (Figure 4 and S Table 11). This pathway uses L-arginine as a carbon source and degrades it to succinate for the TCA. Thus, at 4 hours, the concentrations of L-arginine, GABA -the intermediate molecule in the polyamine putrescine degradation, and succinate had a significant drop of its concentration by 5-fold in both intracellular and extracellular environments remained slightly lower towards the entry of the stationary phase. This was also consistent with a significant transient upregulation of all *puu* operon genes at 4 hours. The expression levels of *puuR*, the main repressor that regulates the intracellular putrescine concentrations by repressing several genes of putrescine utilization and transport, had the highest expression at 4 h. Glutamate and GABA are involved in the acid resistance system 2 (AR2), which enteric bacteria use to survive acidic conditions [19]. A significant increase in the expression of *gadA* and the glutamate decarboxylase at 4 hours suggests that bacteria initially convert L-glutamate to GABA, which is then subsequently degraded since levels of glutamate, GABA, and succinate all rapidly decrease intracellularly at 4 hours compared to 3 hours.

The degradation of putrescine can also form other polyamines such as spermidine and spermine through L-methionine degradation. Spermine was found in large concentrations at 3h and drastically decreased over time. The *S*-Adenosyl-L-methionine, synthesized from the essential amino acid L-methionine was found in low concentration at 4h and 5h (only in E1777) and subsequently converted by the spermidine synthase (encoded by *speE*) to spermidine and later spermine, which was found in lower concentration in the extracellular environment. Other acetylated polyamines derived from spermidine such as *N*_1_-acetylspermidine and *N*_1_, *N*_8_ diacetylspermidine were detected but not significantly changed over time. They displayed mixed patterns with increasing concentrations in both supernatant and intracellular fractions. Acetylation converts the polyamines to a physiologically inert form to protect against polyamine toxicity and is mainly excreted from the cell. On the other hand, spermidine can conjugate with thiol glutathione which plays a role in detoxifying xenobiotics and reactive oxygen species. This thiol was detected at increased concentrations over time intracellularly.

## Discussion

Growth in Luria Bertani (LB) broth is commonly used to analyze bacterial properties. Although most natural habitats of prokaryotes do not resemble the nutrients in LB, the transition to more nutrient-depleted conditions, *i.e*., the stationary phase when growth ceases, is a common phenomenon in nature. We analyzed the transcriptome of two ETEC strains during the transition from late exponential phase/early stationary phase to stationary phase to get a comprehensive view of the regulatory and metabolic pathways involved in ETEC and *E. coli* growth in LB. Our results indicated several distinct steps during the transition into the stationary phase in support of previous studies [20]. Specifically, we identified an interesting transient phase at OD_600_ = 3 at the immediate onset of stationary phase characterized by decreased gene expression of genes involved in iron uptake and more than 10-fold upregulation of operons involved in, *e.g*., dipeptide transport, fucose, and putrescine utilization and indole production. Furthermore, integration of the transcriptome with metabolome analyses highlighted the L-arginine, 4-aminobutanoate, and putrescine degradation pathways forcefully induced at the onset of the stationary phase.

Our results suggested that the transient phase when ETEC/*E. coli* is preparing for stationary phase is characterized by a temporal reduction in iron uptake. Interestingly, a similar response has been reported in the transition phase between the log and stationary phase for *Helicobacter pylori* [21], suggesting this phase alteration and its characteristics occur in several bacterial species. Metal ions such as iron are essential for bacteria but, at the same time, extremely toxic. Iron ions in oxidation state II (ferrous (Fe^2+^) iron are more bioavailable than ferric (Fe^3+^) ions in oxidation state III but more toxic since they may form hydroxyl radicals through the Fenton reaction. Our results demonstrated that, e.g., *fecA* and *fhuA* related to iron uptake and transport of ferrous ion and ferric ion were transiently downregulated during entry into stationary phase. This may have implications in ensuring iron homeostasis in an iron-deficient environment such as the gastrointestinal tract [22–24].

Transcriptome analysis also indicated increased fucose metabolism at the onset of stationary phase at 4h followed by down-regulated at 5h. Fucose is an abundant mucus-derived metabolite in the intestine generated by commensal bacteria [25]. The most common commensal bacterium in the gut is *Bacteroides thetaiotaomicron*, and some *E. coli* can utilize fucose as a carbon source [26]. We confirmed that E2265 used in this study could metabolize L-fucose in a phenotypic assay, but interestingly this trait was not conserved over all ETEC lineages. Several studies have shown the important role of fucose in virulence. For example, in *Salmonella* Typhimurium, *fucI* was significantly upregulated one day after infection in germ-free mice colonized by *Bacteroides thetatiotaomicron*, and the respective mutant had decreased competitiveness *in vivo* [27]. Another example of fucose modulating the virulence is EHEC, which harbors a pathogenicity island LEE containing a two-complement-system (TCS) capable of sensing fucose and transcribing the pathogenicity island LEE [28]. In EHEC, it has also been demonstrated that fucose is important for colonization, suggesting that bacterial pathogens take advantage of using unexploited sugars by commensal bacteria [29]. Hence the induction of fucose utilization might promote colonization of the mucosa of bacteria in the early stationary growth phase. The fact that E2265 belongs to the commonly isolated and globally spread clonal lineage 5 [4] might implicate that fucose metabolism is important for virulence in certain ETEC lineages.

Pathway enrichment analysis identified two significant pathways; degradation of aminobutyrate (GABA) degradation and degradation of 5-hydroxytryptamine (serotonin), to be significantly enriched in cluster III characterized by genes that progressively increased during entry into stationary phase (Fig 2). Genes involved, *e.g., puuE*, *gabD, aldAB, feaB*, and *prr* increased their expression levels up to 10-fold at 4 hours compared to 3 hours and increased further in the last sampling point at 5 hours.

The aminobutyrate degradation pathway (Fig 4) is linked to L-arginine and putrescine degradation pathways. Putrescine and its downstream metabolite spermidine are polyamines present in the gut and introduced by food, microbial, or intestinal cell metabolism [30]. They regulate cellular function in both prokaryotic and eukaryotic cells. The transient phase was characterized by upregulation of the *puu*-regulon. Putrescine is generated by decarboxylation of ornithine or decarboxylation of arginine into agmatine. *E. coli* uses specific importers such as PotFGHI and YdcU, members of the ATP-binding cassette (ABC) transporter family, and PuuP to take up putrescine across the cell membrane. Once putrescine is imported, PuuA γ-glutamylates putrescine resulting in a γ-glutamyl-γ-aminobutyraldehyde, which is oxidized by PuuB and subsequently dehydrogenated by PuuC. Then, PuuD hydrolyzes the resultant γ-glutamyl group, generating and releasing simultaneously γ-aminobutyrate (GABA) and glutamate. PuuE deaminates GABA to succinic semialdehyde, which is oxidated by GabD (which does not belong to the Puu operon) to produce succinic acid that goes to the TCA metabolism [31–33].

In terms of virulence, polyamines, specifically, putrescine has been shown to be actively involved. For instance, a mutant of *potD* in *Streptococcus pneumoniae* displayed attenuated virulence. In *V. cholerae*, defective biofilm formation was observed after deletion of the homologous genes of *E. coli potD*. On the contrary, polyamines might prevent the colonization of the small intestine by *V. cholerae* since high concentrations of polyamines disrupt pili-pili interaction during autoaggregation [34]. In *Salmonella enterica* serovar Typhimurium, exogenous putrescine and spermidine are sensed to prime intracellular survival and induce virulence [35].

Interestingly, the downstream metabolite of putrescine γ-aminobutyric acid (GABA) is a well-known neurotransmitter used as food supplementation in animal husbandry to reduce aggressive behavior and stress [36]. Supplementation of GABA also induces sIgA secretion and increases IL-4 and IL-17 in piglets challenged with porcine ETEC [37]. The AraC-like transcription factor GadX is an activator of the glutamate decarboxylase *gadAB* that converts L-glutamate to GABA. GadX is also a repressor of the transcription factor that activates the expression of adhesion factor bundle forming pilus (*bfp*) and intimin in enteropathogenic *E. coli* [38, 39]. Hence factors that increase GABA might also downregulate virulence factors, and the corresponding rapid degradation of GABA at the transient phase could promote virulence and downregulated immune responses to ETEC. We could, however, not see the corresponding pattern in this study since adhesion factors CS5 and CS6 were not changed. Previous studies on transcriptomes in ETEC have indicated that toxin and CF genes are down-regulated upon binding to cells as well as influenced by bile salts present in the small intestine [11, 40, 41]. We have also previously shown that *eltAB* expression decreases from exponential to stationary growth phase [8].

Tryptophan is cleaved to indole, pyruvate, and NH_4_^+^ by the tryptophanase (TnaA) enzyme expressed by certain bacteria, including *E. coli, Bacteroides* and *Lactobacilli*. Indole is an extracellular signaling molecule well known for affecting different aspects of bacterial physiology, including biofilm formation, in a concentration-dependent manner [42–45]. Indole is also an interspecies signaling molecule. In *E. coli*, the “indole peak” has been described in several studies as a short window of time at entry into the stationary phase where intracellular levels of indole rapidly peak and then decrease again [42]. We found strong induction of *tnaA* expression at 4h compared to 3h and 5h and concomitant rapid decrease of intracellular L-tryptophan at 4h, supporting that the transient phase identified in this study is the same as the indole peak.

Tryptophan metabolism in the gut is important since tryptophan metabolites include indole and neurotransmitters and immunomodulators, including serotonin, tryptamine, and kynurenine. The synthesis of these latter molecules is performed in gut cells like enterochromaffin cells from diet-derived tryptophan and constitutes an important part of the gut-brain axis of neurotransmitters [46]. In this study, metabolomics identified L-5-hydroxytryptophan (5-HTP), the precursor of serotonin (5-hydroxytryptamine), and we also found L-kynurenine (Supplementary Table 6). To our knowledge, only one other study reported that *E. coli* could produce the neurotransmitter serotonin [47]. We performed an additional verification analysis (S Table 12) to search for serotonin in ETEC but were not able to detect it in our samples.

This study supports that commensal and pathogenic bacteria can both degrade and produce neurotransmitters and their intermediate molecules and hence ETEC infection might interact with gut cell signaling and influence the gut-brain axis to a larger extent than previously thought.

In summary, our data suggest that the entry into the stationary phase is a distinct growth phase that might pose ETEC into a stage of increased survival, virulence, and host competitiveness due to lack of need to sequester iron, retained virulence gene expression, and capacity to compete with the commensal flora for host-derived carbon and nitrogen sources such as fucose and putrescine. This study provides a framework for further studies on ETEC gene regulation and comprehensive characterization of transcriptional responses during the transition to the stationary phase that also applies to other bacteria.

## Materials and Methods

### Strains, growth conditions, and bacterial enumeration

The ETEC strains E1777 and E2265 (LT STh/CS5+CS6), both isolated from adult patients with watery diarrhea in Dhaka Bangladesh in 2005 and 2006, respectively, were used in this study [12, 48]. The whole-genome sequences of the two strains are available [18], including a complete assembled chromosome and two plasmids of 142 and 78 kbp, respectively, for E2265 [11, 48]. Bacteria from frozen stock vials were grown on blood plates, and 10 colonies were picked and grown under shaking conditions in 10 ml of LB medium to OD_600_ = 0.8 (10^9^ bacteria/ml) to be used as a starting culture. The starting culture was diluted 100-fold in 20 ml LB medium in a 250 ml Erlenmeyer flask and grown aerated at 150 rpm rotation at 37 °C. Samples for optical density, colony-forming units (cfu), and RNA extraction and metabolomics were withdrawn after 3, 4, and 5 hours. For bacterial enumeration, the track dilution method was performed as described previously [49]. In brief, a 20 μl of bacterial culture was collected every hour up to 6 h and overnight time point and subjected to ten-fold serial dilution in a 96-well plate filled with 180 μl of phosphate-buffered saline (PBS) 1X. 10 μl from each dilution were spotted in a column onto LB agar plates, and the plate was tipped onto its side to allow migration of the spots across the agar surface. This step was performed in duplicate. LB plates were incubated overnight at 37°C, and cfu per ml was quantified by multiplying the number of colonies of each tract by their respective dilution factor and inoculated volume (0.01 ml).

For RNA extraction, bacterial samples from 3, 4, and 5 h time points were immediately mixed with 2 x volume of RNAProtect^®^ (Qiagen) using the manufacturer’s protocol. Samples were stored at −80°C until extraction.

### RNA preparation

Total RNA was prepared from lysozyme and proteinase K lysed bacteria using the RNeasy^®^ Mini Kit (Qiagen, Hilden, Germany) and the instructions provided by the manufacturer for RNA extraction from Gram-negative bacteria. An extra step to remove contaminating DNA on-column was included using the RNase-Free DNase Set (Qiagen). The integrity of the RNA and absence of contaminating DNA was checked by agarose gel electrophoresis, and the RNA concentration was measured spectrophotometrically using a NanoDrop^®^ ND-1000 (NanoDrop Technologies, Wilmington, DE). The RNA was carefully precipitated, washed, and shipped under 99.5% EtOH. The integrity, quality, and concentrations of the RNA were rechecked upon arrival at the sequencing facility at Beijing Genome Institute (BGI), Shenzhen, China, using an Agilent. The RIN values were above 9.8 for all samples.

### RNA-seq

The RNA samples were depleted from rRNA by RiboZero, and Illumina libraries were generated using the TruSeq protocol described by the manufacturer. The libraries were sequenced using the Hi-Seq 200 using a read length of 100 bp. Reads were assembled using the software SOAPdenovo (http://soap.genomics.org.cn/soapdenovo.html). CAP3 assembled all the unigenes from different samples to form a single set of non-redundant unigenes. All unigene sequences were blasted against protein databases using blastx (e-value<0.00001) in the following order: Nr SwissProt:KEGG:COG. Unigene sequences with hits in the first or second database did not go to the next search round against later databases. Then blast results were used to extract CDS from Unigene sequences and translate them into peptide sequences. Blast results information was also used to train ESTScan [50]. CDS of unigenes with no-hit in the blast were predicted by ESTScan and then translated into peptide sequences.

Functional annotations of Unigenes included protein sequence similarity, KEGG Pathway, COG, and Gene Ontology (GO) was performed. First, all-Unigene sequences were searched against protein databases (Nr SwissProt KEGG COG) using blastx (e-value<0.00001). Then, the Blast2GO program [51] was used to get GO annotations of the Unigenes. After getting GO annotation for every Unigene, we used WEGO software [52] to do GO functional classification for all Unigenes.

Each time point’s differential expression was determined using the DESeq2 package (v1.22.1) and R-3.6.0 [53]. The counts were normalized, and the fold change and the log2 of the fold-change were calculated based on the following comparisons: control (3 h) vs. 4 h and control vs. 5 h. Significant genes from each comparison and each strain were filtrated using the following threshold: Padj < 0.05, log2Foldchange < −2 and log2foldchange > 2. PCA and sample distance heatmaps were plotted to visualize the cluster of groups and outliers. Heatmaps of differential genes were generated using the R package *pheatmap* [54]. For temporal gene expression pattern analysis, the Short time-series Expression miner (STEM) was applied as described by Ernst, Nau [17]. For gene ontology (GO) enrichment analysis for biological processes and metabolic pathways, PATHER Overrepresenaton Test (http://www.pantherdb.org/) [55] with an FDR correction applied to all reported P values for the statistical tests.

#### Untargeted Metabolomics

##### Bacterial sampling

The bacterial sampling for metabolomics was performed as described previously [56] with some modifications. A total of 4 ml of bacterial culture were collected and split into two sterile Eppendorf tubes to sample intracellular and secreted metabolites at every time point. In addition, another 100 μl were collected for measurement of the optical density and pH. For extracellular metabolites, 2 ml of bacterial culture was pelleted by centrifugation at 12,000 x *g* for 3 min a tabletop centrifuge, and the supernatants were carefully removed and transferred to a new sterile Eppendorf tube for snap-frizzing in liquid nitrogen. Snap frozen samples were stored at −80°C. The fast filtration method was applied for intracellular metabolites using a 3-place EZ-FitTM Manifold (Millipore®) connected to a single vacuum that supports simultaneous filtration of three samples. A sterile 22 mm diameter MF-Millipore® membrane filter with 0.45-μm pore size was placed onto each manifold, pre-washed with pre-warmed LB medium, and the vacuum set to 50 mbar. 2 ml of the bacterial culture were pipetted in the middle of the filter and subsequently perfused with 5 ml of pre-warmed washing buffer (M9 medium [Sigma-Aldrich] adjusted to pH 7.3) was perfused. Immediately after, the cell-loaded filter was removed and transferred to an Eppendorf tube for snap-freezing in liquid nitrogen. 2 ml aliquots of LB and M9 minimal medium (Sigma-Aldrich) were collected and span-frizzed and used as negative controls. All tubes were kept at −80°C and shipped in dry ice to the Science for Life Laboratory at Uppsala University for metabolite extraction and UPLC-MS analysis.

##### Metabolite Extraction

Supernatants were extracted by the addition of 4 mL of 60:40 ethanol: water solution. The mixture was kept at 78 °C for 3 min, with vigorous mixing every minute. The filters containing the bacterial pellet were transferred to a 4 ml solution and kept at 78 °C for 3 min, with vigorous mixing every minute. The samples were transferred to Eppendorf tubes on ice and centrifuged at 13500 rpm for 5 min, at 4 °C. The supernatant was collected and dried under vacuum on a Speedvac concentrator. The pellet was re-dissolved and injected onto the UPLC-MS system.

##### UPLC-MS Analysis

Ultra-high-performance liquid chromatography coupled to a mass spectrometer (UPLC-MS) was used to identify metabolites, which differ between the three different time points. Mass spectrometric analysis was performed on an Acquity UPLC system connected to a Synapt G2 Q-TOF mass spectrometer, both from Waters Corporation (Milford, MA, USA). The system was controlled using the MassLynx software package v 4.1, also from Waters. The separation was performed on an Acquity UPLC® HSS T3 column (1.8 μm, 100 × 2.1 mm) from Waters Corporation. The mobile phase consisted of 0.1% formic acid in MilliQ water (A) and 0.1% formic acid in LC-MS grade methanol (B). The column temperature was 40 °C and the mobile phase gradient applied was as follows: 0-2 min, 0% B; 2-15 min, 0-100 % B; 15-18 min, 100 % B; 18-20 min, 100-0 % B; 20-25 min, 0 % B, with a flow rate of 0.3 ml/min.

The samples were introduced into the q-TOF using positive electrospray ionization. The capillary voltage was set to 2.50 kV and the cone voltage was 40 V. The source temperature was 100 °C, the cone gas flow 50 l/min, and the desolvation gas flow 600 l/h. The instrument was operated in MSE mode, the scan range was m/z = 50-1200, and the scan time was 0.3 s. A solution of sodium formate (0.5 mM in 2-propanol: water, 90:10, v/v) was used to calibrate the instrument, and a solution of leucine-encephalin (2 ng/μl in acetonitrile: 0.1% formic acid in the water, 50:50, v/v) was used for the lock mass correction at an injection rate of 30 s.

##### Data analysis

The obtained UPLC-MS data comparing the different time points were analyzed using the XCMS software package under R (version 3.3.0) to perform peak detection, alignment, peak filling, and integration. The peaks were annotated by comparing their m/z values to the exact molecular masses of all the online platform Metaboanalyst for pathway analysis. The *Escherichia coli* K-12 MG1655 from the KEGG database was used for metabolite identification. Confirmed metabolites were co-injected with the bacterial samples for the highest level of confirmation. The structures for the significantly altered metabolites were validated with authentic internal standards, as detailed in Figure S12. PCA and heatmaps were performed in R software. The metabolite-metabolite correlations were computed using the R function *rcorr* from the Hmisc package (https://cran.r-project.org/web/packages/Hmisc/index.html). A multiple parametric statistic t-test was performed in GraphPad to compare the means of two paired groups, *i.e*., 3 h vs. 4 h, and multiple comparison corrections were applied using the Holm-Šídák method. *P-value* < 0.05 was set as the threshold for significance.

## Acknowledgments

The study was supported by the Swedish Research Council (dnr 2011-2435, dnr 2014-02639, dnr 2017-01812, and dnr 2020-01941), VINNOVA (2011-03491), and the Swedish Foundation for Strategic Research, SSF (SB12-0072) to ÅS. This study was also supported by the Swedish Research Council (dnr 2016-04423) and a generous start-up grant from the Science for Life Laboratory to DG. I.N. is partially supported by the National Institute of General Medical Sciences of the National Institutes of Health (award P20GM125503). Finally, the authors wish to express their gratitude to the Beijing Genome Institute (BGI) staff, Shenzhen, China, for Illumina sequencing.

## Supplementary Figures

**S Figure 1.**
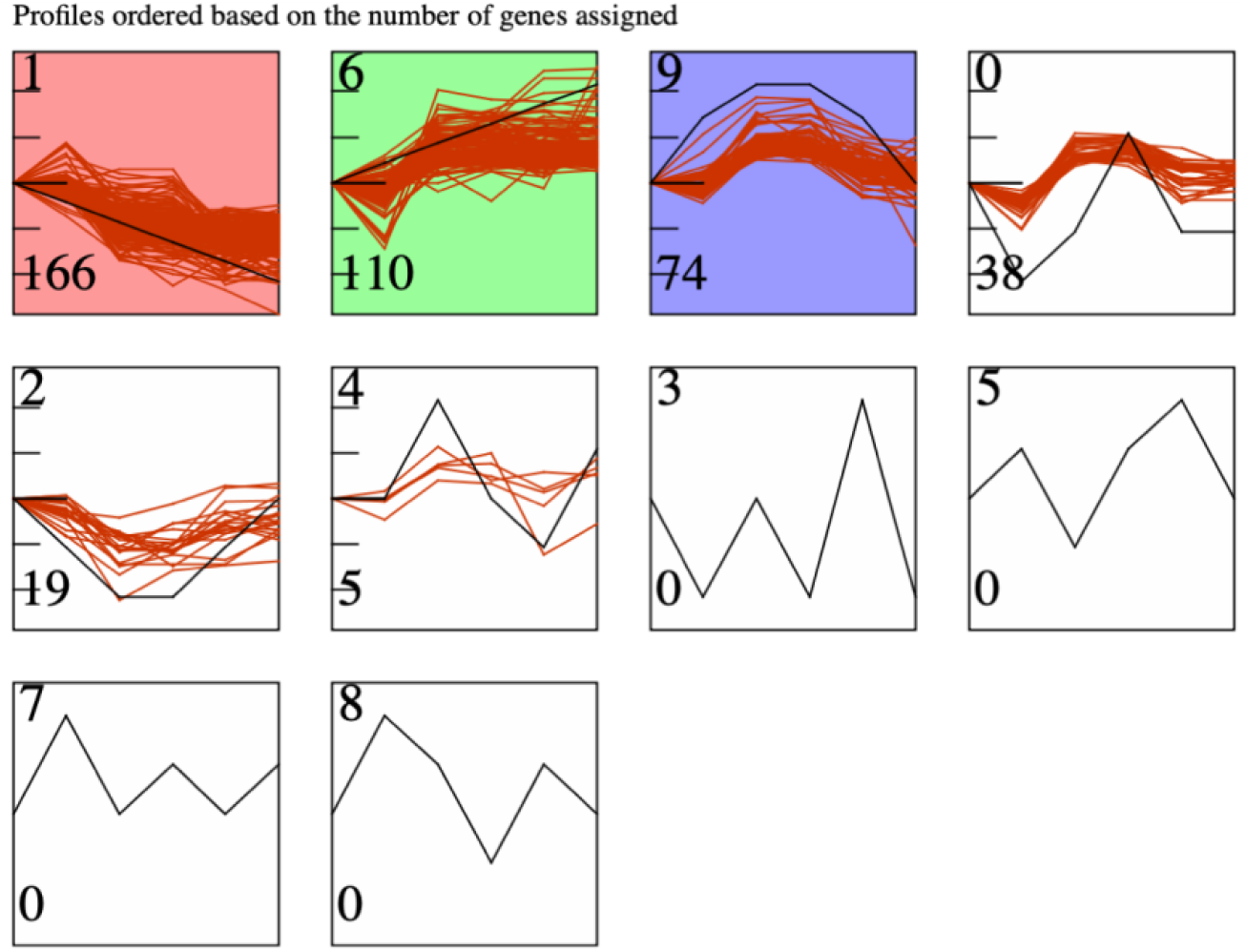
Idenified gene expression patterns by STEM. The profile number on the top left corner of each profile box was assigned by STEM and the number on the bottom left represents the number of genes included in the cluster. Clusters with a *p*-value >0.05 were colored.

**S Figure 2.**
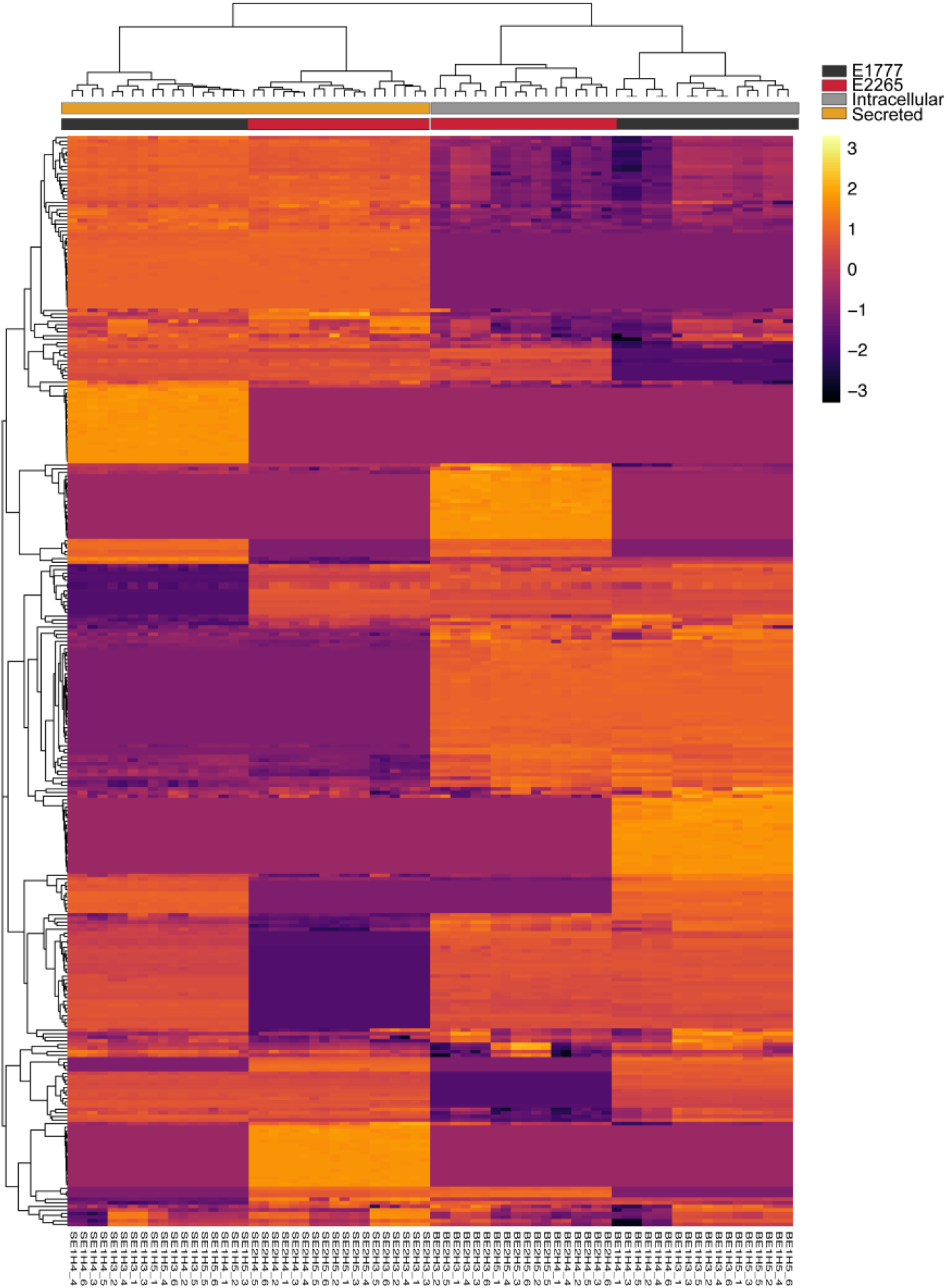
Heatmap representation of the abundance of 288 intracellular and secreted metabolites detected in E2265 and E1777.

**S Figure 3.**
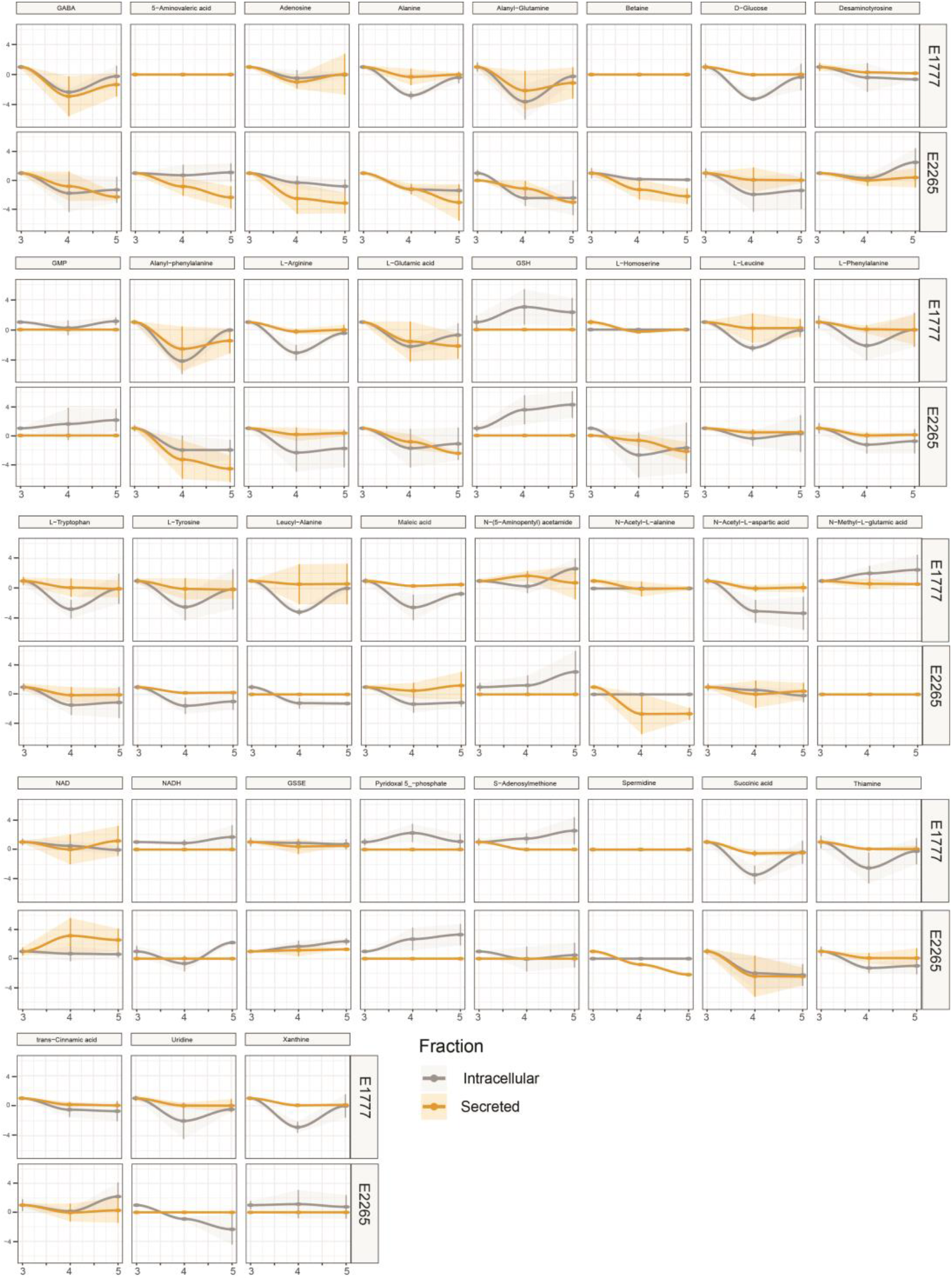
Data of the concentration of all significant metabolites are represented in the serial charts. The metabolite concentration is adjusted to 3 h, and the data is presented in fold-change±SD.

